# Molecular mechanisms of re-emerging chloramphenicol susceptibility in extended-spectrum beta-lactamase producing Enterobacterales

**DOI:** 10.1101/2023.11.16.567242

**Authors:** Fabrice E Graf, Richard N Goodman, Sarah Gallichan, Sally Forrest, Esther Picton-Barlow, Alice J Fraser, Minh-Duy Phan, Madalitso Mphasa, Alasdair T M Hubbard, Patrick Musicha, Mark A Schembri, Adam P Roberts, Thomas Edwards, Joseph M Lewis, Nicholas A Feasey

## Abstract

Infections with Enterobacterales (E) are increasingly difficult to treat due to antimicrobial resistance. After ceftriaxone replaced chloramphenicol (CHL) as empiric therapy for suspected sepsis in Malawi in 2004, ESBL-E rapidly emerged. Concurrently, resistance to CHL in *Escherichia coli* and *Klebsiella* spp. decreased, raising the possibility of CHL re-introduction. However, many phenotypically susceptible isolates still carry CHL acetyltransferase (CAT) genes.

We used a combination of genomics, phenotypic susceptibility assays, experimental evolution and functional assays for CAT activity to understand the molecular mechanisms and stability of this re-emerging CHL susceptibility.

Of 840 Malawian isolates, 31% had discordant CHL susceptibility genotype-phenotype, and we selected 42 isolates for in-depth analysis. Stable degradation of *cat* genes by insertion sequences led to re-emergence of CHL susceptibility. Our study suggests CHL could be reintroduced as reserve agent for critically ill patients with ESBL-E infections in Malawi and similar settings and highlights the ongoing challenges in inferring antimicrobial resistance from sequence data.

## Introduction

Antimicrobial resistance (AMR) is a major threat to global health. Drug resistant bacterial infections were estimated to be associated with 4.95 million deaths in 2019, with sub-Saharan Africa the worst affected region [1]. Among the most problematic drug resistant bacteria are extended-spectrum beta-lactamase (ESBL) producing Enterobacterales (E) which are resistant to 3^rd^-generation cephalosporins (3GC) and classified as priority pathogens by the WHO [2]. ESBL-E infections are associated with higher morbidity, mortality and economic burden in the treatment of bloodstream infections in Malawi [3, 4], and in low resource settings such as Malawi access to effective treatment alternatives (e.g. carbapenems, or beta-lactam/beta-lactamase inhibitor combinations) is often lacking. The prevalence of ESBL-E rapidly increased in Malawi after ceftriaxone replaced chloramphenicol (CHL) as a first-line empiric therapy for suspected sepsis from 2004 [5, 6].

CHL is an inhibitor of protein synthesis and broad-spectrum antibiotic discovered in 1947 [7], which is now rarely used, primarily because of the risk of severe adverse effects [8]. These include dose-unrelated aplastic anaemia, which is irreversible and fatal [9] but rare (incidence between 1:19,000 – 1:270,000 [10]), dose-related, reversible bone marrow suppression [11] and Grey Baby Syndrome [12]. As effective and safer broad-spectrum beta-lactam antibiotics were introduced, they replaced CHL in most settings. The dominant mechanism of CHL resistance is mediated by CHL acetyltransferases (CAT) inactivating CHL by acetylation [13]. Other mechanisms include efflux pumps, inactivation by phosphotransferases, target site mutations and decreased membrane permeability [13].

While resistance to 3GCs, fluoroquinolones and aminoglycosides increased in Malawi, resistance to CHL decreased as its use declined [5]. *Escherichia coli* and *Klebsiella pneumoniae* populations showed decreasing proportions of CHL resistant isolates, from around 80% in 1998-2004 to around 50% and below from 2012, sparking an interest in whether CHL can be re-introduced as a treatment option for ESBL-E infections in critically ill patients where there is no alternative therapy [5, 14]. Some bacteria isolated in Malawi, however, have a discordant CHL genotype-phenotype, i.e. they still harbour CHL resistance genes despite being phenotypically susceptible [15–18]. Therefore, simple CHL resistance gene loss caused, for example, by lineage or plasmid replacement in the bacterial population is unlikely. With the prospect of using CHL as a potential reserve agent for 3GC resistant infections it is important to understand (i) the molecular mechanism(s) of this re-emerging CHL susceptibility, i.e., the molecular basis of CHL susceptibility genotype-phenotype mismatches, (ii) the stability of the phenotypic susceptibility and (iii) how widespread this phenomenon is. Here, we investigated a collection of ESBL *E. coli* and *K. pneumoniae* with CHL susceptibility genotype-phenotype mismatches. Using functional assays in combination with genomic data, we determined the CHL sensitivities, CAT enzyme activity, and functional expression of *cat* genes, to understand the molecular mechanism of re-emerging CHL susceptibility. Further, we tested the stability of CHL susceptibility in several isolates employing experimental evolution with increasing concentrations of CHL and used co-occurrence analysis of AMR genes and investigated the spread of *cat* alleles in the context of sequence types to identify potential drivers to explain the high proportion of CHL susceptibility in ESBL-E populations in Malawi.

## Results

### Genotype-phenotype mismatch for chloramphenicol resistance

We screened a collection of 566 *E. coli* and 274 *K. pneumoniae* species complex (*KpSC*) isolates previously isolated from patients admitted to Queen Elizabeth Central Hospital (QECH) in Blantyre, Malawi. All isolates had been whole-genome sequenced, of which 164 (93 *E. coli*, 71 *KpSC*) were from sentinel surveillance of bacteraemia [5, 15, 16] and 676 isolates (473 *E. coli*, 203 *KpSC*) were asymptomatically carried ESBL-isolates from stool, collected in a research study [14, 17, 18] (Supplementary Table 1). Of the total isolates, 53.5% (449/840) were phenotypically susceptible to CHL (Fig. 1a) and 31.0% (260/840) had a discordant genotype-phenotype. 45.0% of phenotypically susceptible isolates (202/449) harboured CHL resistance genes; the majority were chloramphenicol acetyltransferase (*cat*) genes (193/202) (Fig. 1b). The most prevalent *cat* genes were *catB4* (229) followed by *catA1* (189), *catA2* (104) and *catB3* (19) (Fig. 1c). Other known CHL resistance genes, *cmlA1* (29)*, cmlA5* (23) and *floR* (14), were less common in the collection. We selected a subset of 42 isolates, 27 *E. coli* (13 CHL-R, 14 CHL-S) and 15 *K. pneumoniae* (6 CHL-R, 9 CHL-S), based on a genotype-phenotype mismatch and control isolates (same sequence type (ST) if available) for further in-depth functional analysis (Table 1) to investigate the molecular mechanism of those mismatches. First, we determined the CHL susceptibility for each isolate by broth microdilution and compared the result to previously collected AST data [5, 14], determined for CHL using the disc diffusion method. For 81.0 % (34/42) isolates, microbroth dilution confirmed the result of disc testing and we concluded that phenotype-genotype mismatches were not explained by inaccurate phenotypic data. Five of the eight isolates that had discordant disc diffusion and MIC results were within 2x of the breakpoint for CHL (>8 µg/mL) Table1. We used the more sensitive MIC result to classify isolates as resistant or susceptible.

**Fig. 1.**
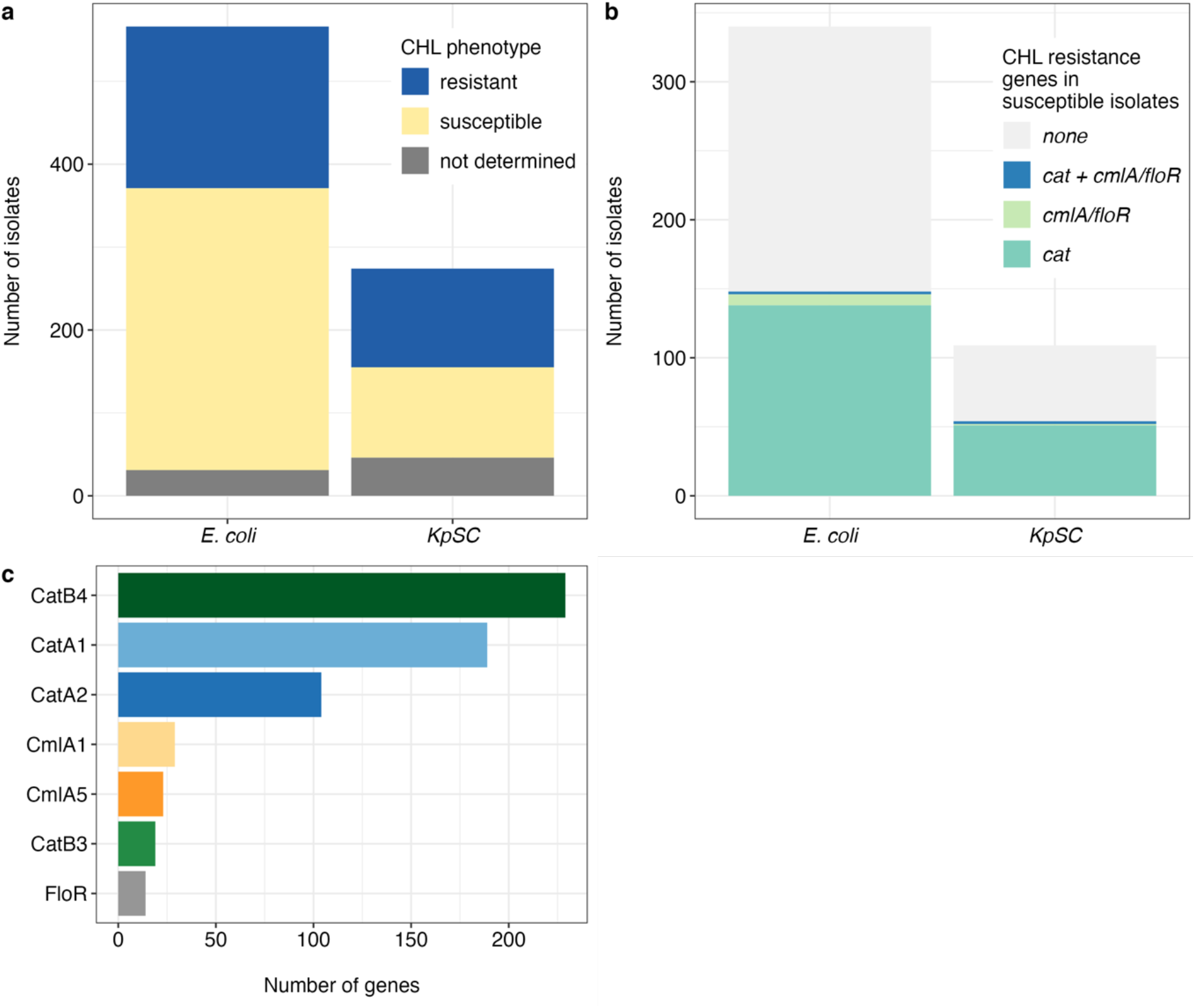
Characteristics of isolates. **a** Chloramphenicol susceptibility of *E. coli* and *KpSC* isolates. **b** Number of CHL susceptible isolates carrying *cat* genes, *floR* or *cmlA* genes, both (*cat* + *floR* or *cmlA*) or no CHL resistance genes. **c** Number of CHL resistance genes present in the 840 Malawian isolates.

**Table 1.** Isolates functionally characterised in this study.

| ID | ST | Species | CHL MIC | R/S | CHL R gene(s) present |
| --- | --- | --- | --- | --- | --- |
| CAE137 | 648 | *E. coli* | 512 | R | *catA1*, *catB4* |
| CAD110 | 424 | *E. coli* | 512 | R | *catA2*, *catB4*, *cmlA5* |
| CAB199 | 44 | *E. coli* | 128 | R | *catB4*, *floR* |
| CAC122 | 101 | *E. coli* | 32 | R | *cat_pC221*, *cmlA1* |
| CAC10A | 167 | *E. coli* | 16 | R | *catA1* |
| CAG10A | 46 | *E. coli* | 16 | R | *catB3* |
| CAI104 | 46 | *E. coli* | 16 | R | *catB3* |
| CAD10B | 46 | *E. coli* | 16 | R | *catB3* |
| CAD105 | 46 | *E. coli* | 16 | R | *catB3* |
| CAC10H | 617 | *E. coli* | 16 | R | *catB4* |
| CAN108 | 617 | *E. coli* | 16 | R | none |
| CAB17W | 131 | *E. coli* | 16 | R | none |
| CAE11U | 167 | *E. coli* | 12 | R | none |
| CAB184 | 44 | *E. coli* | 8 | S | *catA1, catB4* |
| CAI10X | 167 | *E. coli* | 8 | S | *catA1* |
| CAI10E | 167 | *E. coli* | 8 | S | *catA1* |
| CAC124 | 131 | *E. coli* | 8 | S | *catB4* |
| CAD13N | 410 | *E. coli* | 8 | S | *catB4* |
| CAE12L | 44 | *E. coli* | 8 | S | *catB4* |
| CAD13L | 410 | *E. coli* | 8 | S | *catB4* |
| CAC116 | 167 | *E. coli* | 8 | S | none |
| CAC10W | 10 | *E. coli* | 8 | S | none |
| CAE12U | 10 | *E. coli* | 8 | S | none |
| CAI10Z | Novel | *E. coli* | 4 | S | *catA1, catB4* |
| CAM10K | 443 | *E. coli* | 4 | S | *catA1* |
| CAJ10A | 131 | *E. coli* | 4 | S | none |
| CAC13M | 44 | *E. coli* | 4 | S | none |
| CAF11G | 15 | *KpSC* | 512 | R | *catA1, catB4, cmlA5* |
| CAC11N | 45 | *KpSC* | >512 | R | *catA2, catB4, cmlA5* |
| CAE12L | 340 | *KpSC* | >512 | R | *catA2, cmlA5* |
| CAE135 | 340 | *KpSC* | >512 | R | *catA2, cmlA5* |
| CAB15M | 152 | *KpSC* | 256 | R | *catA1, catB4* |
| CAC13Y | 307 | *KpSC* | 64 | R | *catB4* |
| CAF102 | 15 | *KpSC* | 8 | S | *catB4* |
| CAC14G | 1427 | *KpSC* | 8 | S | *catB4* |
| CAB16G | 14 | *KpSC* | 4 | S | *catB4* |
| CAC105 | 307 | *KpSC* | 4 | S | *catB4* |
| CAC13D | 340 | *KpSC* | 4 | S | *catB4* |
| CAG19R | 1427 | *KpSC* | 4 | S | *catB4* |
| CAD137 | 14 | *KpSC* | 4 | S | none |
| CAL11H | 307 | *KpSC* | 4 | S | none |
| CAD107 | 14 | *KpSC* | 2 | S | none |
| MG1655 | - | *E. coli* | >512 | R | pEB1-*catA1* |
| MG1655 | - | *E. coli* | >512 | R | pEB1-*catA2* |
| MG1655 | - | *E. coli* | >512 | R | pEB1-*catB3* |
| MG1655 | - | *E. coli* | 8 | S | pEB1-*catB4* |
| MG1655 | - | *E. coli* | 8 | S | none |

### No CAT activity in susceptible isolates

Next, we tested all 42 isolates for CAT enzyme activity to determine whether *cat* genes were functionally expressed. We adapted the rapid CAT assay (rCAT) [19] as an indirect measure of enzyme activity. The rCAT assay measures free sulfhydryl groups of the CAT substrate and acetyl donor acetyl-coenzyme A in cell lysates and a colour change to yellow indicating CAT enzyme activity can be spectrophotometrically measured.

All phenotypically CHL-S isolates, irrespective of the presence of a *cat* gene, were negative for CAT activity (Fig. 2). All but five resistant isolates (CAC10A, CAC122, CAC10H, CAB119, CAC13Y) with a *cat* gene showed CAT activity. Three CHL-R/rCAT negative isolates carried *catB4* genes and this was the only *cat* gene present. Isolate CAB199 (CHL MIC = 128 µg/mL) co-carried a *floR* gene which could contribute to the CHL-R phenotype. Isolates CAC13Y (CHL MIC = 64 µg/mL) and CAC10H (CHL MIC = 16 µg/mL) had no other detectable CHL resistance determinants whereas CAC10A had a *catA1* but a weak CHL-R phenotype (MIC = 16 µg/mL). We used the disc-diffusion (dCAT) assay [20] as a secondary and independent assay, which is based on the ability of CAT producing bacteria to cross-protect susceptible bacteria by inactivating CHL. Tested isolates (CAB119, CAC13Y) failed to cross-protect a sensitive strain, thus confirming that *catB4* in those isolates is non-functional (supplementary Fig. 1). One isolate (CAC122) that carried *cat_pC221*, a *cat* gene originating from *Staphylococcus aureus* [21], did not show enzyme activity and its resistance phenotype is likely explained by *cmlA1*. This *cat* gene was shown to be inducible and regulated through translational attenuation [22]; we thus tested this isolate with sub-MIC concentration of CHL in culture media but no CAT activity was observed. We concluded that resistant isolates with *catA1*, *catA2* and *catB3* have functional CATs whereas isolates with *catB4* do not. Phenotypically susceptible isolates with or without *cat* genes did not show CAT activity demonstrating that *cat* genes in those CHL-S isolates are not expressed.

**Fig. 2.**
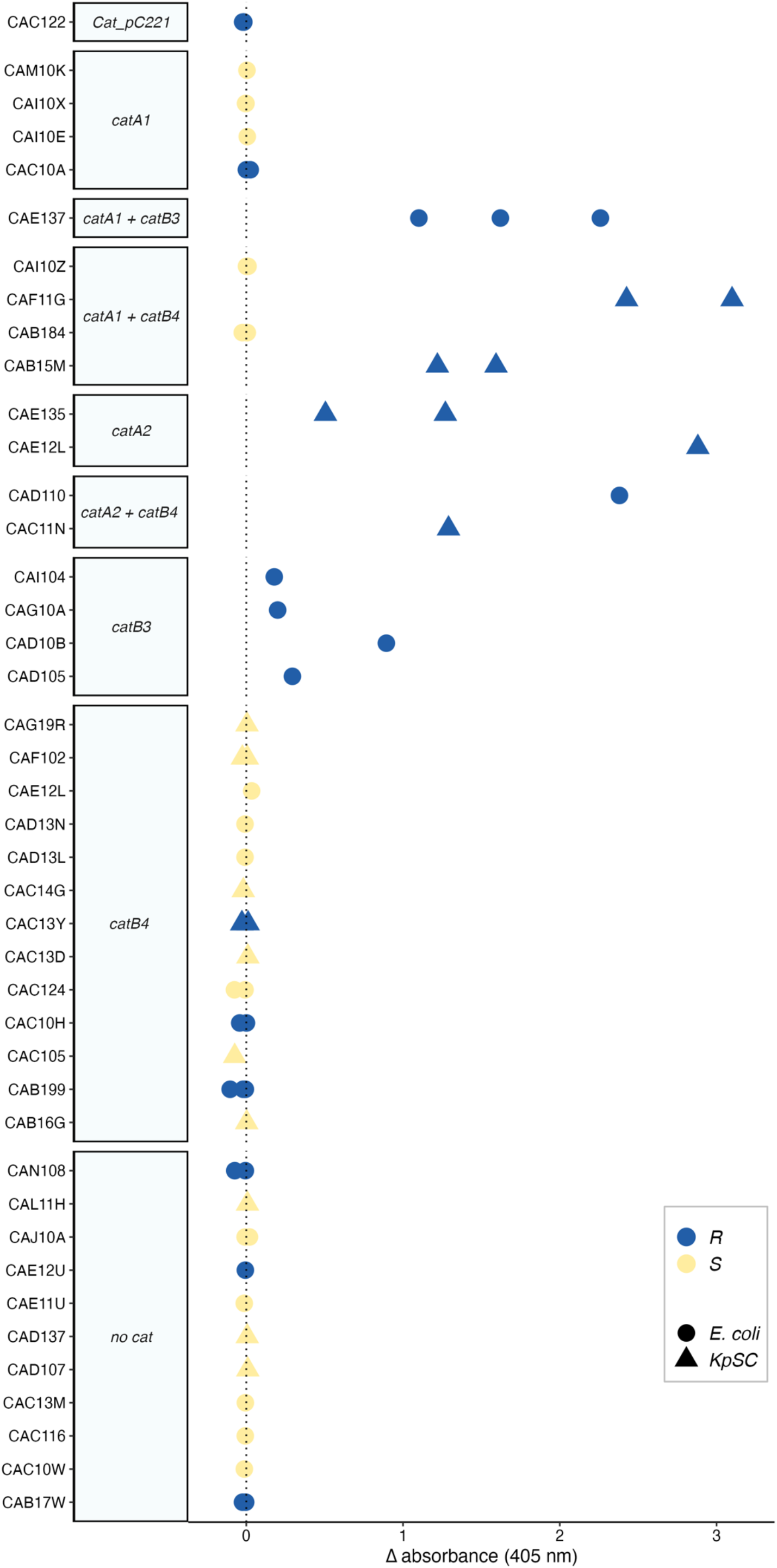
CAT enzyme activity. CAT enzyme activity was measured using the rapid CAT assay. The difference ((signal + CHL) - (signal - CHL) for a single isolate (n = 2-6)) in absorbance at 405 nm is given for each of the 42 isolates. Colour indicates if the isolate is phenotypically resistant (R, blue) or susceptible (S, yellow) to chloramphenicol based on microbroth dilution and the shape indicates *E. coli* (circle) or *K. pneumoniae* (triangle). The Y axis is ordered according to presence of *cat* genes.

### Functional expression of *catB4* does not confer CHL resistance

To assess the effect of the genetic context of *cat* genes in our isolates, we cloned the coding sequence (CDS) of the dominant *cat* variants present in the collection (*catA1*, *catA2*, *catB3* and *catB4*) into the expression vector pEB1-*sfGFP* [23] under the constitutive *proC* promoter, by replacing *sfGFP*, and tested their susceptibility to CHL in a clean genetic background of *E. coli* MG1655. MG1655 pEB1-*catB4* had an MIC of 8 µg/mL identical to the MG1655 pEB1-sGFP control whereas all other *cat* variants had an MIC of >512 µg/mL confirming that *catB4* is a non-functional variant (Table 1).

### *catB4* is a non-functional truncated *catB3*

Functional CAT assays (rCAT and dCAT) demonstrated that *catB4* in the genetic context of tested isolates was non-functional and ectopic expression of *catB4* confirmed that the CDS does not produce a functional product that confers CHL resistance. We previously observed that in addition to a full length assembled *catB4* (549 bp) many isolates had several partial assemblies in their genomes of approximately 107 bp (Supplementary Fig. 2a, [24]). We investigated the 107 bp (position 443-549 of *catB4*) by BLASTn against NCBI’s nucleotide collection and obtained many hits among different Enterobacterales with 100% query cover and sequence identity matching the “IS*6*-like element IS*26* family transposase”. We next compared *catB4* with *catB3* by pairwise sequence alignment. Both CDS share 100% sequence identity for the first 442 bases and only differ after position 443, corresponding to the last 107 bp of *catB4* matching with the IS*26* sequences (Fig. 3a). All but seven *catB4* assemblies in our isolate collection share 100% sequence identity (Supplementary Fig. 2b) with *catB3* but only until position 442 bp (full length *catB3* is 633 bp). Therefore, we concluded that *catB4* is a truncated variant of *catB3* and suggest it should be called *catB3Δ^443-633^*. IS*26*-mediated deletions of adjacent DNA have previously been reported [25, 26].

**Fig. 3.**
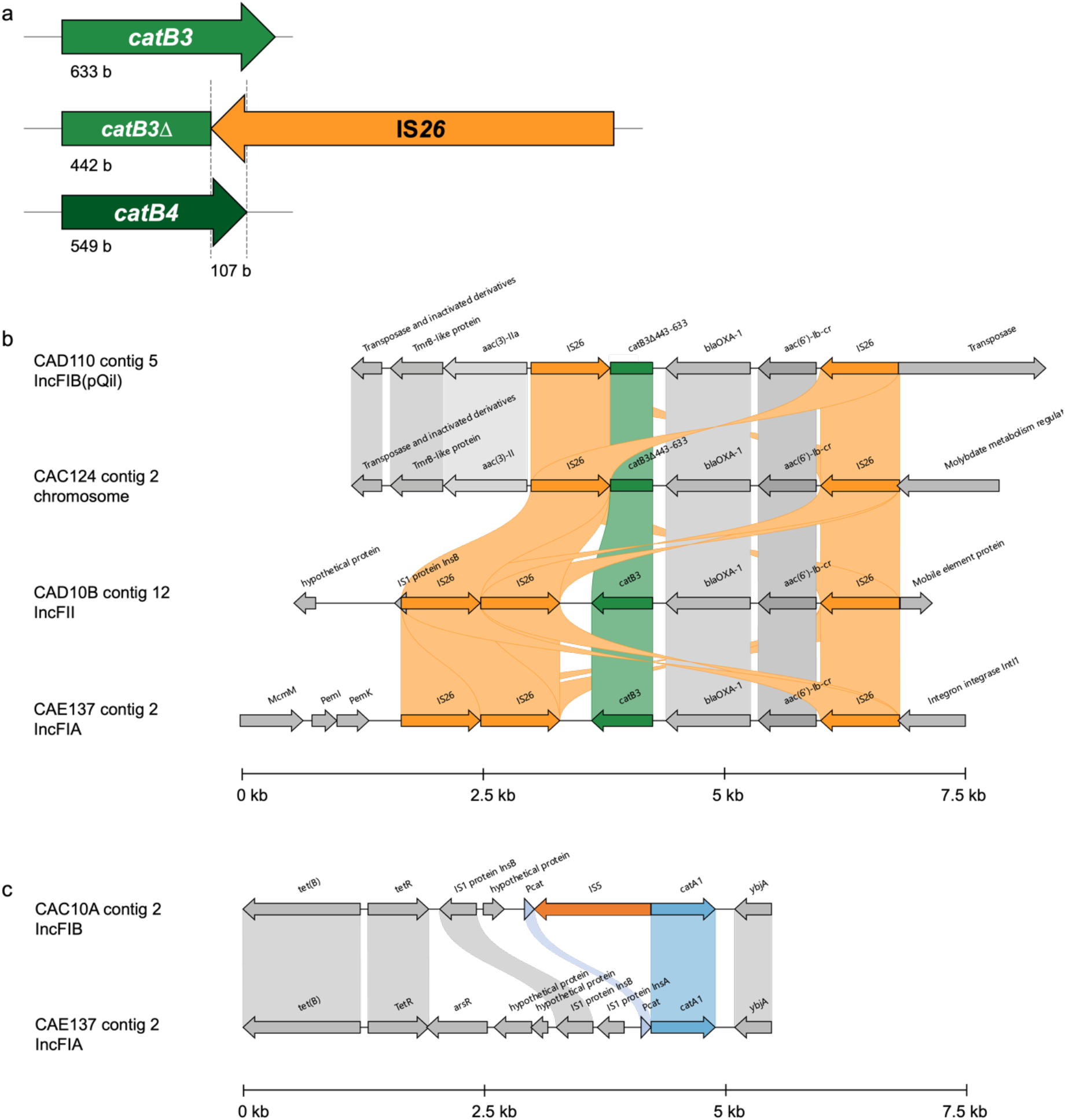
*Cat* gene degradation by insertion sequences. **a** Schematic of IS*26* truncation of *catB3*. **b** Alignment of four contigs from four different isolates containing either *catB3* and *catB3Δ^443-633^*. **c** Alignment of two genomes containing *catA1*, with and without an IS*5* insertion into the *catA1* promoter.

We traced *catB4* back in the literature and found it was first described, to the best of our knowledge, in two plasmids (pEK499 and pEK516) [27]. Both plasmids had been moved into different *E. coli* lab strains and did not confer CHL resistance, consistent with our data showing that *catB4* is not a functional CAT. Because this had been annotated as *catB4* variant in the ARG-ANNOT (and SRST2) databases [28] we tested two additional databases for AMR genes. Both, the CARD database [29] and ResFinder [30] called the truncated *catB3* correctly.

We selected five *E. coli* isolates for long-read sequencing to investigate the genomic context of *catB3Δ^443-633^* (Supplementary Table 3). Aligning four closed contigs containing *catB3* and its truncated variant showed a similar and common genetic feature of *aac(6’)-Ib-cr* - *bla*_OXA-1_ - *catB3* flanked by two IS*26* elements. There was an additional IS*26* downstream of the wild type *catB3* gene (Fig. 3b). In the four isolates this feature is located on a different replicon, on the chromosome and three different IncF plasmids suggestive of moving as transposable element independent of a single plasmid-type. As this genetic feature has previously been shown to be part of an integron cassette [31, 32], we ran IntegronFinder and found *attC* sites upstream of *bla*_OXA-1_ and downstream of *catB3*, however, the truncation of *catB3* removed the *attC* site, likely precluding its ability to move between or within integron(s). Further, the integrase gene is missing in all but one isolate (CAE137) where it is interrupted by IS*26* (Fig. 3b).

### IS*5* insertion into *catA1* promoter causes reversal of CHL resistance

Of 9/42 isolates with a *catA1*, only four had a CHL resistance phenotype consistent with its genotype (Fig. 2). We therefore PCR-probed these nine isolates with primers specific for *catA1* to confirm its presence. Three isolates containing *catA1* that were CHL-R had the expected amplicon size of 150 bp, however three CHL-S isolates and CAC10A had a larger amplicon than expected of ∼1400 bp, (supplementary Fig. 3). We assembled the genomes of isolates with *catA1* and aligned the contigs containing *catA1*. Two contigs (from isolates CAI10X & CAI10E) showed a presumed insertion of 1199 bp, starting 6 bp upstream of the CDS, matching the larger than expected PCR amplicon and BLASTn revealed a 100% match to an IS*5* element. In addition, we purified and Sanger-sequenced the ∼1400 bp PCR product and confirmed the insert as an IS*5* element in three of the isolates. Long-read sequencing of isolate CAC10A confirmed the insertion of IS*5* as a single element into the promoter region of *catA1* (Fig. 3c). Overall, of the isolates with *catA1,* 3/9 were CHL-R with detectible CAT activity (Fig. 2) and had an expected *catA1* amplicon size (150bp); isolate CAC10A was CHL-R with an MIC close to the breakpoint (16 mg/L), no detectable CAT activity and IS*5* inserted into the *catA1* promoter region. The remainder (5/9) were CHL-S, without detectable activity on the rCAT assay; in 3/5 this was likely mediated by IS*5* insertion interfering with transcription. We adjusted primers from a previously developed high-resolution melting (HRM) assay [33] to capture IS*5-catA1* and the truncated *catB3Δ^443-633^*, and demonstrated that they can be clearly distinguished from the wild type genes (Supplementary Fig. 4), highlighting the potential of this molecular diagnostic test for CHL resistance to discriminate between functional and non-functional *cat* genes.

### Degradation of *cat* genes by insertion sequences is stable

IS*5* insertion into the promoter of *catA1* and truncation of *catB3* are the main mechanisms for the observed re-emerging CHL susceptibility in our collection of isolates. To test if these mutation and insertion events could be reversed and potentially result in a rapid re-emergence of *cat* expression and CHL resistance, we experimentally evolved three CHL susceptible isolates: CAI10Z (*catA1*, *catB3Δ^443-633^*), CAM10K (*catA1*) and CAI10X (*catA1*) in LB broth with increasing concentrations of CHL for 7 days (n=3 per isolate). An equal number of replicates were evolved in LB without selection. All evolved populations under CHL selection grew until 64 µg/mL CHL (8-16 -fold increase in MIC). Increasing incubation for another 24 h enabled recovery of some populations in 128 µg/mL (i.e., clearly visible growth) but none of the CHL populations grew in 256 µg/mL CHL (Supplementary Fig. 5a). All control populations evolved in LB grew until the experiment was stopped at day 7. Since we were interested if CHL pressure can result in a re-activation of *cat* genes we tested all evolved populations with the rCAT assay (LB evolved from day 7, CHL evolved last surviving population, day 5, day 6 or day 7). All populations examined tested negative on the rCAT assay (Supplementary Fig. 5b) and PCR-probing of evolved strains showed that the IS*5* insertion was still present upstream of *catA1* (supplementary Fig. 5c), confirming that other mutations rather than (re-)expression of *catA1* are responsible for the resistance phenotype.

### The genomic locus of *catB3Δ^443-633^* is conserved and widespread

The truncated *catB3Δ^443-633^* is common among our isolates and seems to be highly conserved in the genomic context of *aac(6’)-Ib-cr - bla*_OXA-1_ - *catB3*. To investigate potential drivers, we ran a co-occurrence analysis across 772 genomes in our collection from Malawi (68 of the 840 total isolates failed assembly quality check) (Fig. S6, supplementary Table 4). We applied the probabilistic model from [34] to determine whether each pair of genes had an observed co-occurrence which was significantly different from the expected co-occurrence. The co-occurrence relationships across all AMR genes with at least one other gene is depicted in Fig. S6. Both, *catB3* and *catB3Δ^443-633^*had a positive co-occurrence relationship with *bla*_OXA-1_ and *aac(6’)-Ib-cr* and the beta-lactamase genes *bla*_CTX-M-15_ and *bla*_TEM-1B_. The *catB3* and *catB3Δ^443-633^* genes had a negative co-occurrence as they were never found to co-occur on the same genome (Fig. 4).

**Fig. 4.**
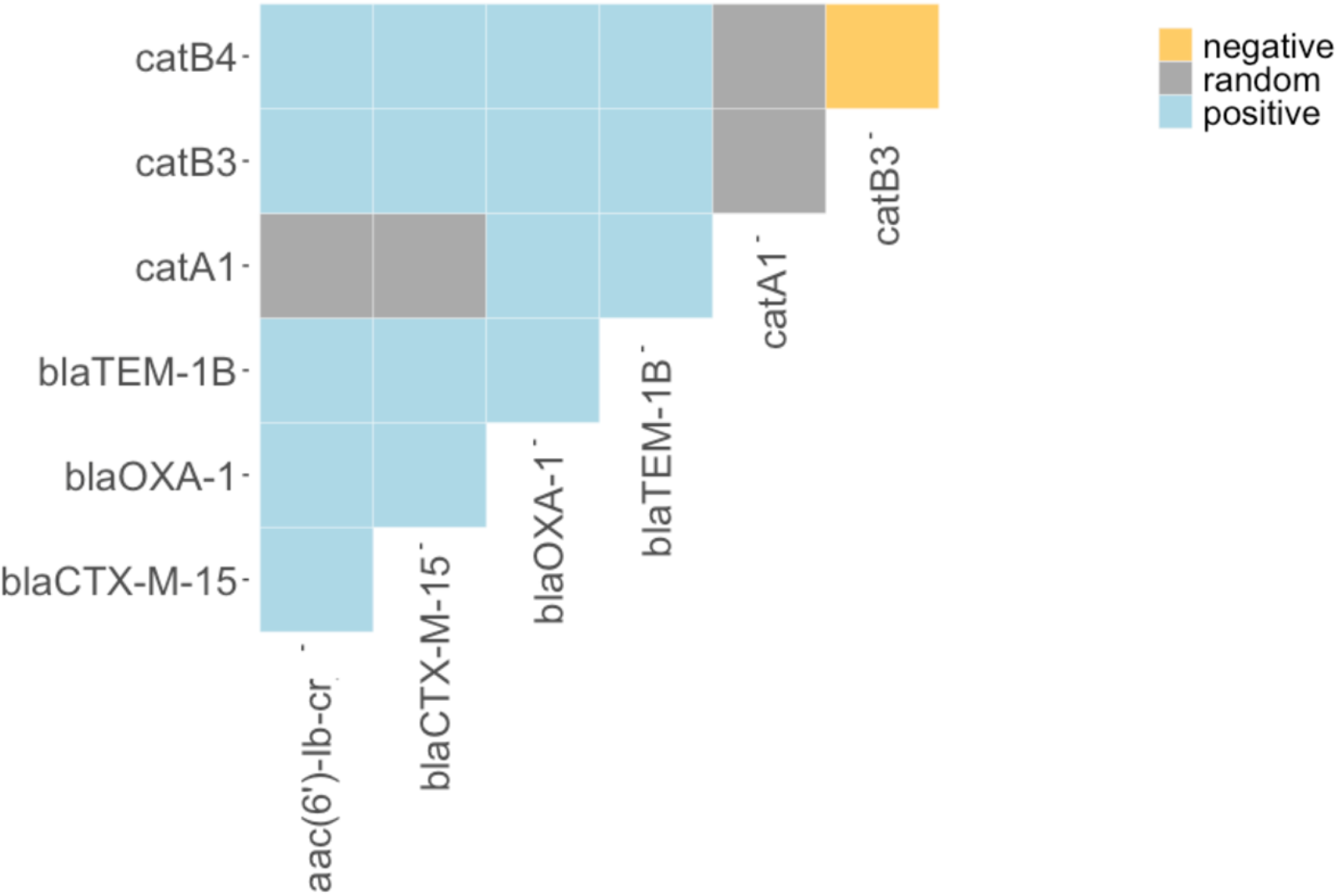
Co-occurrence networks of AMR genes. Heatmap displaying co-occurrence relationships between AMR genes as either positive (blue), random (grey) or negative (orange). These are probabilistic values based on the difference in expected and observed frequencies of co-occurrence between each pair of genes, these values were obtained by applying the probabilistic model from [34]. Co-occurrence across select genes including *cat genes*, *aac(6’)-Ib-cr, bla*_OXA-1_, bla*_CTX-M-15_* and *bla_TEM-1_*.

In our isolates, *catB3Δ^443-633^* is much more common than *catB3*, and thus we investigated if this pseudogene is restricted to Malawi or common elsewhere. We queried “catB3” in NCBI‘s Microbial Browser for Identification of Genetic and Genomic Elements (Micro-BIGG-E) [35] and investigated 46,667 isolates (data download: 4/8/2023) with an annotated *catB3* gene. Strikingly, 30,819 (66%) showed a coverage to the reference of 70%, which corresponds to the IS*26* truncated variant described here (Supplementary Fig. 7a). We tried to trace back the emergence of this truncation. From 2006 onwards (only including years with > 100 isolates) the truncated variant was dominant over the wild type, and we could not conclude when it emerged (Supplementary Fig. 7b). *Klebsiella* spp., *E. coli* and *Enterobacter* spp. Dominantly show the truncated gene whereas *Salmonella*, *Acinetobacter* and *Pseudomonas*, typically harbour wild type *catB3* (Supplementary Fig. 7c). Differentiating by host where the isolates have been collected is heavily biased towards humans where the dominant truncated variant is much more common. This is also seen in companion animals (cats and dogs) whereas in most food animals, the wild type proportion is close to 100%, perhaps indicative of ongoing selection for chloramphenicol resistance through use of phenicols in veterinary medicine and agriculture (supplementary Fig. 7d). Most geographic locations show higher proportions of the truncated variant with a few exceptions in China and Australia (Supplementary Fig. 7e). Next, we investigated the spread of *catB3Δ^443-633^* in the context of ST to potentially link clonal expansion and *catB3Δ^443-633^* carried by these clones *in E. coli*. The *catB3Δ^443-633^* gene was restricted to six STs (410, 44, 648, 405, 617, 131) in our Malawi isolates (Fig. 5), and five of those STs are among the top 8 most common STs in our collection (Supplementary Fig. 8). We expanded our analysis and looked at a collection of 10k genomes consisting of the top 100 common STs in *E. coli* (previously curated and downloaded from Enterobase (2020)[36] where each ST was randomly sampled to select 100 genomes. All six STs with *catB3Δ^443-633^* in Malawi also dominantly harbour this *cat* gene in this extended genome collection (Fig. 5). In phylogroup B2 (predominantly human isolates associated with urinary tract and bloodstream infections) *catB3Δ^443-633^* is common in ST131 and to a lesser extend ST1193 (Fig. 5), both of which represent recently emerged globally dominant antibiotic resistant clones [37, 38]. The ubiquitousness of *catB3Δ^443-633^* may thus be linked to its association with distinct and successful lineages.

**Fig. 5.**
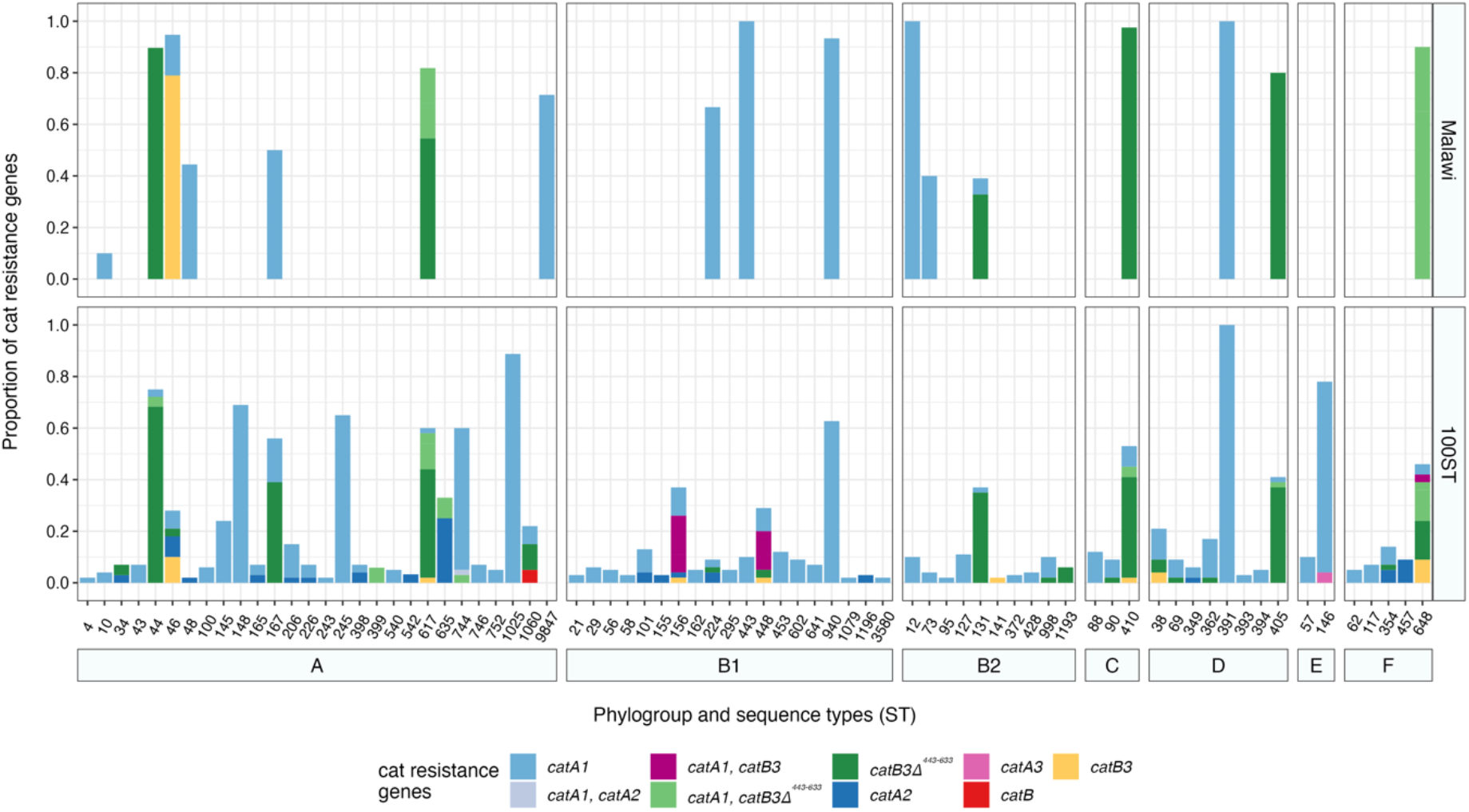
Cat genes occurrence and proportions per Sequence type in *E. coli* isolates from Malawi included in our study (top panel) and in a 10k genome collection of 100 randomly selected genomes of the top 100 most common STs from *E. coli*.

## Discussion

In Malawi, as in much of sub-Saharan Africa, the rapid spread of ESBL-E coupled with the scarcity of Watch and Reserve antibiotics (i.e. carbapenems) has rendered many severe bacterial infections untreatable, therefore the re-emergence of chloramphenicol susceptibility is potentially important. 34.2% (1666/4874) of non-salmonella Enterobacterales were phenotypically CHL susceptible in bloodstream infections in Malawi from 1998–2016 [5]. Here, we analysed a collection of phenotypically and genotypically characterised *E. coli* and *KpSC* from Malawi to understand the molecular basis of CHL-S genotype-phenotype mismatches.

Since many phenotypically susceptible isolates in our collection carried *cat* genes, we first investigated if they were still functional and expressed. None of the CHL-S isolates we tested had functional CATs and we found that *cat* gene interruption was caused by insertion sequences. This is in contrast to a previous study reporting an *E. coli* strain being susceptible to CHL despite a functional CAT [39] which was later attributed to low level of acetyl-coenzyme A linked to mutations in efflux pumps [40]. Combined functional analysis of *catB4* with a CAT enzyme assays, expression in a clean genetic background as well as genomic investigation confirmed that *catB4* is non-functional and is in fact a *catB3* that has been truncated by an IS*26* element. In other cases, an IS*5* element has integrated into the promoter region of *catA1* and likely interfered with transcription. In the latter case, calling AMR genes from all databases with sequence data will yield a wild type gene and currently classify the isolate as resistant since the mutation is outside of the CDS. The truncated *catB3* will be called correctly if the annotation of *catB4* is removed in the ARG-ANNOT/SRST2 databases, which we strongly support.

To effectively re-introduce CHL as a reserve treatment option for Enterobacterales infections confirmed as 3GC-resistant, we must consider the potential for reversion to CHL-resistance. Isolates carrying IS*5*-*catA1* could potentially rapidly revert to high-level resistance since the *catA1* CDS is still present in the genome. Our data using experimental evolution suggest that this is not the case and selection with CHL did not lead to expression of *catA1.* One of the evolved isolates co-carried a truncated *catB3* and we expected no reversion to a functional *catB3* upon CHL selection because the missing 3’ prime end of the gene is no longer present in the genome. Indeed, no functional *cat* emerged.

A decrease in CHL resistant isolates has also been reported for *Salmonella* Typhimurium, which was caused via loss of CHL resistance genes by lineage replacement [41]. Our study adds to a developing evidence base for CHL in treatment of MDR Gram-negative pathogens; several studies have reported high or increasing rates of phenotypic CHL susceptibility among MDR Gram-negative bacteria suggesting CHL as a viable alternative treatment option in those settings [42–44]. Further, CHL is an affordable and useful antimicrobial in terms of bioavailability, tissue penetration and broad spectrum of action [45, 46]. However, the prevalence of CHL susceptibility and the rare but severe side-effects of CHL, preclude the use of CHL for empirical management of sepsis in our setting: we instead envisage CHL to be used in critically ill patients with confirmed ESBL-E and CHL-S infection as a reserve agent. This has the added advantage of keeping selection pressure for CHL resistance low but does require rapid determination of CHL susceptibility phenotype.

In the Malawian isolates the gold-standard AST correctly determined phenotypic CHL susceptibility, however, rapid molecular diagnostics have the potential to be faster and low cost. We applied the HRM assay to distinguish between the *cat* mutants found in our study. Co-occurrence analysis showed that *catB3* and *catB3Δ^443-633^* nearly always co-occurred with *aac(6’)-Ib-cr* and *bla*_OXA-1_. This conserved feature of *aac(6’)-Ib-cr*-*bla*_OXA-1_-*catB* flanked by IS*26*, which was confirmed by long-read sequencing, has previously been associated with an integron [32] and was found to contain *attC* sites when *catB3* was present. However, the truncation by IS*26* removed one of the *attC* sites. This, in combination with loss of the integrase gene, likely led to functional loss of mobilisation of genes within the integron and the high conservation of this locus is confirmatory. The association of *catB3Δ^443-633^*with *bla*_CTX-M-15_, the most common ESBL gene found in *E. coli* isolates worldwide [47] and among our Malawian isolates [17, 18], may point towards the co-location of these genes on a plasmid, though a larger selection of isolates will need to be (long-read) sequenced to determine the spread of such a plasmid. Selection for 3GC resistance and co-occurring *bla*_CTX-M-15_ with *catB3Δ^443-633^* could have led to a genetic hitchhiking of the latter which could explain the high levels of CHL susceptible isolates among ESBLs. However, our isolate collection is heavily biased towards ESBL and may thus limit the interpretation of this co-occurrence.

Expanding our analysis of *catB3Δ^443-633^* revealed that this single truncation is globally more common than the wild type gene. It is intriguing that a truncated non-functional gene is so widespread, and we hypothesise this is due to the specific genetic context of a loss of integron activity, and association with other AMR genes as well as with IS*26*. It has previously been shown that IS*26* and pseudo-compound transposons (i.e. IS*26* bounded transposons) move 50x more frequently if there is another IS*26* to target [48] and move by co-integration [49]. This mode of mobility likely enhances persistence. Additionally, association of *catB3Δ^443-633^*with globally dominant lineages of *E. coli* (e.g. ST131) as seen in our ST analysis, could be a likely driver that has contributed to the widespread occurrence of this truncated variant.

Our study highlights that antibiotic susceptibility can re-emerge following a reduction or cessation in use. Plasticity and the high levels of horizontal gene transfer among Enterobacterales can result in degradation of AMR genes by mobile genetic elements which can persist in the absence of selection. Further, the context of the AMR gene determines the phenotypic resistance and database curation is crucial to infer resistance from the genotypes. We strongly recommend integrating functional context to databases and including additional features such as genetic context, including promoters and transcription start site. Many AMR genes only cause clinically relevant resistance when they are moved to locations in the genome where expression is upregulated, such as downstream of a strong promoter [50]. This is problematic with functional metagenomic discovery of AMR genes since the mere presence of an AMR gene is not indicative of phenotypic resistance. Lastly, our data support the reintroduction of CHL as a last-line treatment option for patients critically ill with ESBL-E infections in Malawi and similar settings.

## Materials and methods

### Antimicrobial susceptibility testing

AST was performed using broth microdilution according the EUCAST guidelines (v.4.0) in cation adjusted MullerHinton broth (MH2, 90922, Merck) in duplicates. If results were inconsistent or different to the CHL susceptibility phenotype previously determined with disc diffusion, AST was repeated. Broth microdilution data was used to determine the phenotypic classification of the subset of 42 isolates. Isolate are classified according to EUCAST (v.12.0); isolates with an MIC ≤ 8 µg/mL were classified as susceptible and isolates > 8 µg/mL as resistant. In EUCAST v.13.0, CHL breakpoints are no longer listed, stating a screening cut-off (CHL MIC > 16 µg/mL) can be used to distinguish wild type from acquired resistance.

### Functional CAT assay (rCAT)

We adjusted the rapid CAT assay from [19] to enable read-out in 96-well plates using a spectrophotometer. Cultures of isolates to be tested were set up in replicates in 1-2 mL LB and incubated at 37 °C with 220 rpm. 100 µL of overnight culture was transferred into a microcentrifuge tube containing 500 µL PBS and centrifuge at 8000 rpm for 3’ to wash and pellet cells. The supernatant was completely removed using a pipette, the pellet resuspended in 250 µL lysis buffer (1 M NaCl, 0.01 M EDTA, 0.05 % SDS) and cells lysed while incubating for 1 h at 37 °C with occasionally vortexing the tubes. Next, 50 µL of the lysed cells were transferred into each of 4 wells of a 96-well plate. To each well 50 µL reaction buffer was added (for 1 mL, 400 µL 0.2 M Tris; 400 µL 5 mM acetyl-coA; 200 µL 10 mM DTNB were mixed). DTNB was added last to the wells, since DTNB was reported to inhibit certain CAT enzymes [19]. DTNB was always prepared fresh since stored DTNB (for even 1 day at 4 °C in the dark) increased the background. Acetyl-coA was always prepared on ice and aliquots immediately frozen at -20 °C and used within 2 weeks.

50 µL 5 mM CHL was added to the first two wells and 50 µL ddH_2_O to the wells three and four of each isolate. The plate was incubated for > 10 min at 37 °C - longer incubation led to a higher signal for positive samples while the background did not change - up to 1 hour tested. The plate was read with a plate reader (GloMax® Discover Microplate Reader (Promega) or CLARIOstar Plus Microplate Reader (BMG Labtech)) at absorbance of 405 nm or 412 nm and absorption from lysed cells without CHL was subtracted from the value with CHL, hence, positive values indicate CAT activity, no activity give a value around zero.

### Chemical competent cells and transformation

MG1655 were made chemically competent as follows: A single colony of MG1655 was picked from an LB plate and incubated in 3 mL LB broth on at 37 °C with 220 rpm for 16 hours. The culture was diluted 1:100 (1 mL in 100 mL) in a 500 mL flask and incubated at 37 °C with 220 rpm until OD_600_ reached 0.3 – 0.4. The flask was immediately cooled in an ice-water slurry before the cells were transferred to pre-chilled 50 mL Falcon tubes. Cells were centrifuged at 4000 rpm at 4 °C, supernatant discarded and the cell pellet resuspended in 10 ml ice-cold CaCl_2_ solution (60 mM CaCl_2_, 15% glycerol, 10 mM PIPES pH 7). The centrifugation step was repeated, and resuspension was kept on ice for 30 minutes. After another centrifugation step the cells were resuspended in 1 mL ice-cold CaCl_2_ solution and aliquots of 50 µL or 100 µL were frozen at -70 °C.

For transformation, the chemically competent cells were thawed on ice. 2 µL of plasmid DNA (pEB1-variants; 0.5 ng/mL) was added and incubated on ice for 30 minutes. Cells were heat-shocked for 30 seconds at 42 °C in a water bath and immediately put back on ice for 2 minutes. 900 µL SOC media was added to cells and incubated for 1 hour at 37 °C 220 rpm before being spread on LB kanamycin (50 µg/mL) plates.

### Cloning and functional expression of *cat* genes

Cat gene variants were amplified using primers spanning the coding sequence with 20 bp overhangs homologous to the pEB1-plasmid cloning site (supplementary Table S2). Gibson assembly [51] (NEBuilder® HiFi DNA Assembly Master Mix, E2621S, NEB, UK) was used to clone *catA1*, *catA2*, *catB3* and *catB4* into pEB1. Plasmids were verified by colony PCR and Sanger sequencing of the insert regions using pEB1_sequencing primers (supplementary Table S2). Plasmids were extracted using QIAprep Spin Miniprep Kit (27106, Qiagen) and transformed into competent MG1655.

### Experimental evolution of CHL resistance

Isolates were retrieved from frozen stock on LB agar plates. A single colony was used to inoculate a 1 mL starting culture and incubated at 37 °C with 220 rpm (the ancestor). Then, for each tested isolate the starting culture was diluted 1:100 into 3 tubes with 3 mL LB broth (control populations) and 3 tubes with LB and 0.5x the MIC of CHL (selected populations) of the isolate. All population were incubated for 24 h at 37 °C with 220 rpm and then again diluted 1:100 in fresh LB broth with or without CHL. In the selected population the CHL concentration was doubled with each passage. This was repeated for 7 days or until the population became extinct.

### DNA extraction

DNA was extracted with MasterPure™ Complete DNA & RNA Purification Kit (MC89010, LGC Biosearch Technologies). Isolates were grown overnight on blood agar (PP0120-9090, E&O labs) at 37°C and colonies were picked and washed in 500 μl of sterile PBS (10010023, Fisher Scientific) by centrifugation at 10,000 x g for 3 minutes. Pellets were resuspended in 300 μl Tissue and Cell Lysis Solution supplemented with 1 μl of Proteinase K solution. Tubes were incubated at 65°C shaking at 1000 rpm for 5 minutes. Samples were placed on ice for 3 minutes. 1 μl of RNase A solution was added and the samples incubated at 37°C for 30 minutes.

Samples were placed on ice for 5 minutes then 150 μl of MPC Protein Precipitation Reagent was added and samples vortexed for 10 s. The resulting protein debris was pelleted by centrifugation at 10,000 x g for 10 minutes at 4°C. The supernatant was aspirated and added to 500 μl of isopropanol (15631700, Fisher Scientific) and the tube inverted 30-40 times to precipitate the nucleic acids which were then pelleted by centrifugation at 10,000 x g for 10 minutes at 4°C and supernatant discarded. The pellet was washed twice in 70% ethanol and resuspended in 50 μl of TE Buffer. The quantity of DNA was then measured using Qubit dsDNA Quantitation, Broad Range kit (Q32850, Thermo Scientific) on a Qubit 4 Fluorometer (Thermo Scientific).

### Long-read sequencing

Five *E. coli* from the subset of 42 isolates with a genotype-phenotype mismatch, were re-streaked from glycerol stocks onto antibiotic supplemented LB Agar and incubated at 37°C for 16 h. A single colony was picked and transferred to LB broth supplemented with antibiotics. Antibiotic supplementation included chloramphenicol and/or ampicillin depending on the resistance profile of each strain. Genomic DNA was extracted using the Firemonkey High Molecular DNA Extraction Kit (Revolugen, UK) according to the manufacturers protocol.

The strains were long-read sequenced on a MinION device (Oxford Nanopore Technologies (ONT), UK). Library prep was carried out according to the manufacturer’s protocol (ONT, UK) using the SQK-NBD114.24 Ligation Sequencing and Native Barcoding Kit. The DNA library was quantified at several stages in the library prep using a Qubit Fluorometer (ThermoFisher Scientific, Massachusetts, USA), the TapeStation 4100 (Agilent, California, USA) was used to determine the molar concentration and 10-20 fmol were loaded onto the flow cell. Sequencing was carried out using a FLO-MIN114 (R10.4.1) flow cell (ONT, UK) on a MinION Mk1B sequencer, running for 72 h at a translocation speed of 400bp/s. Data acquisition used the MinKNOW software (v22.08.9).

### Bioinformatic analysis

The raw fast5 files from the MinION sequencing run were basecalled using Guppy v6.4.2 with the super accuracy (sup) model for DNA sequencing on the R10.4.1 flowcell with the E8.2 motor protein and the 400bp/s translocation speed. Nanoplot v1.38.1 [52] was used to check the quality of the sequencing reads and the parameters of the sequencing run. The basecalled fastq files were demultiplexed using the guppy_barcoder from Guppy v6.3.8 with the SQK-NBD114-24 kit. The sequence reads were assembled with flye v2.9.1 [53] using trestle mode. Seqkit v0.15.0 [54] was used to determine basic statistics and contig sizes (supplementary Table 3). The assembled contigs were annotated with RAST [55] and any query MGEs aligned against the ISfinder database [56], Mobile Element Finder database [57] and IntegronFinder [58]. Resfinder was used to determine resistance genes [59]. Annotated assemblies were visualised in Snapgene, and clinker [60] was used on the Genbank files to align the contigs and highlight any homologous genetic clusters.

### Co-occurrence analysis of AMR genes

Genome sequences (all illumina short read data) for our assembled isolate collection were obtained from the European Nucleotide Archive: PRJEB8265, PRJEB28522, PRJEB26677 and PRJEB36486 [15–18]. Fastq files were downloaded in April 2023. Initially, the paired-end short-read fastq files were downloaded and trimmed with cutadapt [61]. All reads were analysed with FASTQC and found to pass over half the quality control determinants, sequences with a Phred quality score less than Q20 across the length of the reads were excluded. Reads were assembled using SPAdes v3.11.1 [62]. 68 genomes failed initial QC or assembly and the dataset taken forward for analysis contained 495 *E. coli* and 277 *KpSC* genomes. The quality of the assembled contigs was assessed using the stat command from Seqkit v0.15.0 [54]. The final dataset contained 772 assembled fasta files.

Abricate v0.0.9 (https://github.com/tseemann/abricate) was used to screen for AMR genes and create output tables, sequences were compared against the Resfinder [59] database at 60% minimum length and 90% percentage identity using the BLASTn algorithm. The R programming language v 4.3.1 was used to convert the abricate output tables into an appropriate format for further data analysis and visualisation, including a binary presence/absence AMR gene table, with the tidyverse (v1.3.0) package [63]. Any annotation of *catB3* with less than 75% coverage was labelled *catB4*. The cooccur package v1.3 [64] was used to create a co-occurrence matrix containing the probabilistic values which represent whether a co-occurrence relationship is observed significantly more or less than could have happened by chance. This used the probabilistic model of co-occurrence [34]. If a pair of genes were observed to co-occur significantly less than expected by chance their relationship was termed negative, if they were observed to co-occur significantly more it was termed positive and any pair of genes which were observed to co-occur with no significant difference from their expected value were termed random. A significant difference was defined as having a p-value ≤ 0.05. Select genes conferring resistance to the phenicol, beta-lactam and aminoglycoside drug classes were then selected from the original presence/absence table for further analysis. Heatmaps to display co-occurrence were visualised using ggplot v 3.4.3 [63] and pheatmap v 1.0.12. Clustering in the heatmap used hierarchical clustering with Euclidian distance and the complete method.

### ST analysis

The genome assemblies of 100 randomly selected isolates from each of the 100 most common STs in Enterobase were downloaded on 18/12/2020. Extra genomes from ST391, ST44, ST940 and ST9847 with release date before 18/12/2020 were added later to match with the STs from the Malawi dataset. The STs of the Malawi dataset were determined using MLST (v2.23.0). The presence of *cat* genes was detected using AMRFinderPlus version 3.11.20 with database version 2023-09-26.1 [65]. The data was analysed using R (v4.1.3) with the tidyverse package (v1.3.1) and plotted using ggplot2 (v3.4.2), ggpubr (v0.4.0), and RColorBrewer (v1.1-2).

## Supplementary materials and data availability

Supplementary Figures 1-8 and supplementary Tables 2 & 3 are accessible in the supplementary material of this manuscript. Reads from all isolates previously sequenced and used in this study are accessible in the European Nucleotide Archive (ENA) under project IDs PRJEB8265, PRJEB26677, PRJEB28522 and PRJEB36486 and ENA accession numbers for isolates linked to metadata are available in the Supplementary Table 1. Long-read sequence data have been submitted to the National Center for Biotechnology Information (NCBI) under the BioProject ID PRJNA1040831, BioSample accession numbers for sequenced isolates are in supplementary Table 3. The R scripts used to generate analyses, figures and tables, and supplementary Tables 1 & 4 are available from the GitHub repository https://github.com/FEGraf/CHL-Malawi.

## Author contributions

The study was conceived by F.E.G., A.P.R., and N.A.F. The methodology was devised by F.E.G., R.N.G., MD.P., M.A.S., A.P.R., T.E., J.M.L. and N.A.F. Investigations were undertaken by F.E.G., R.N.G., S.G., S.F., E.PB., A.J.F., MD.P., M.M., A.T.M.H., P.M., T.E., and J.M.L. Formal analysis was done F.E.G., R.N.G., MD.P., P.M., M.A.S., A.P.R., T.E., J.M.L. and N.A.F. The original draft was prepared by F.E.G., R.N.G, J.M.L. and N.A.F., and then reviewed and edited by all authors. Supervision was by F.E.G., T.E., A.P.R. and N.A.F.

## Acknowledgements

We wish to thank Dr Anne Farewell for *E. coli* MG1655. pEB1-sfGFP was a kind gift from Philippe Cluzel (Addgene: http://n2t.net/addgene:103983). We also wish to thank Dr Simon Wagstaff for the GPU base-calling of Oxford Nanopore data and Dr Nadja Wipf for critically reviewing R-code.

