## Supplementary materials for "Molecular mechanisms of re-emerging chloramphenicol susceptibility in extended-spectrum beta-lactamase producing Enterobacterales"

S Figure 1 – dCAT assay

S Figure 2 – Assemblies and percentage identity of *catB4*.

S Figure 3 – PCR probing of *catA1*

S Figure 4 – HRM assay

S Figure 5 – Stability of IS*5-catA1* upon CHL selection

S Figure 6 – Co-occurrence networks of AMR genes

S Figure 7 – Frequency distribution of *catB3*

S Figure 8 – Number of isolates per sequence type

S Table 1 – Isolates and ENA accession number used in this study; accessible at https://github.com/FEGraf/CHL-Malawi

S Table 2 – Primers used in this study

S Table 3 – Long-read stats

S Table 4 – Co-occurrence table; accessible ay https://github.com/FEGraf/CHL-Malawi

Supplementary methods

Supplementary references

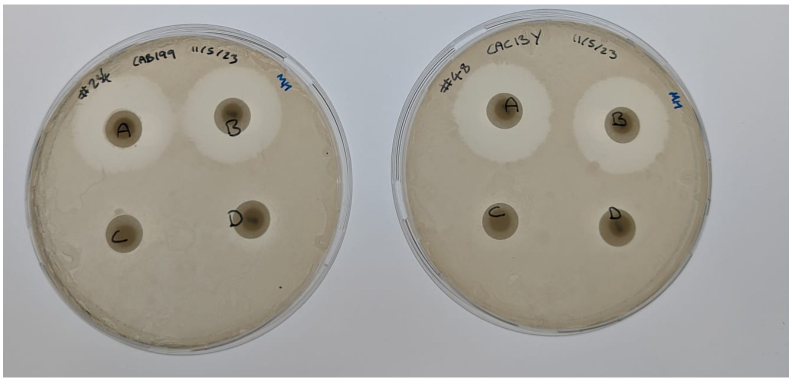

**Supplementary Figure 1 | dCAT assay** measuring cross-protection from CAT enzyme producing isolates. Labels on plate: A tested isolate, B susceptible control, C susceptible - control without chloramphenicol (CHL) disc, D + control (*catA1*), Isolates CAB119 & CAC13Y fail to inactivate CHL from disc.

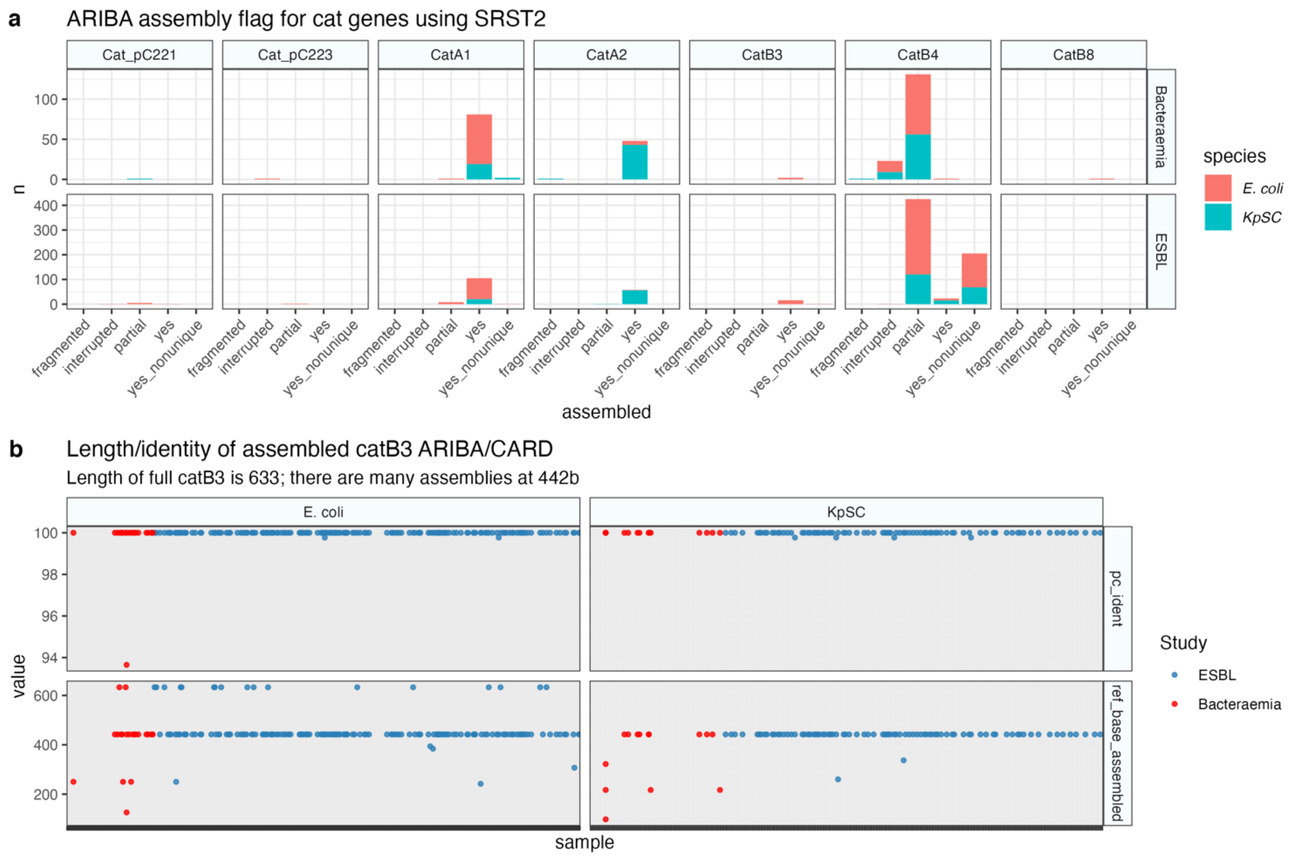


**Supplementary Figure 2 | Assemblies and percentage identity of *catB4* a** assemblies of *cat* genes among the ESBL and the bacteraemia sentinel isolates using Ariba/SRST2. Fully assembled (yes), fully assembled multiple times (yes_nonunique), partially assembled (partial) or interrupted assembly (interrupted). **b** Percentage identity and number of bases assembled to ref of *catB4* compared to *catB3* from the CARD database.

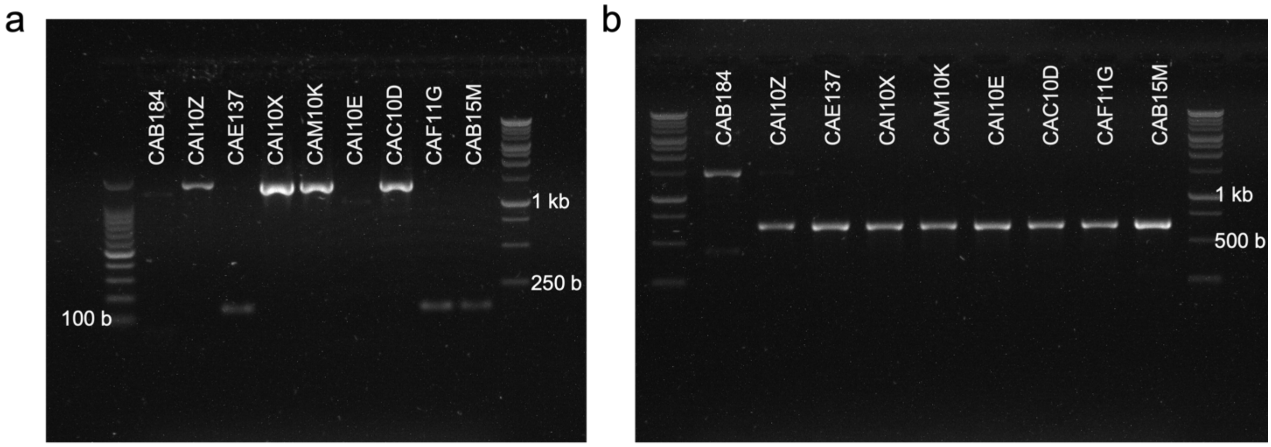


**Supplementary Figure 3 | PCR probing for *catA1*.** PCR amplicon ran on 1% agarose gel: **a** *catA1* probing: expected size 150 bp. Lane 1: 100 b Marker, lane 11: 1kb plus Marker (Promega, UK). **b** catA1-CDS primers: expected amplicon size 660 bp. Lane 1 & 11: 1kb plus Marker (Promega, UK).

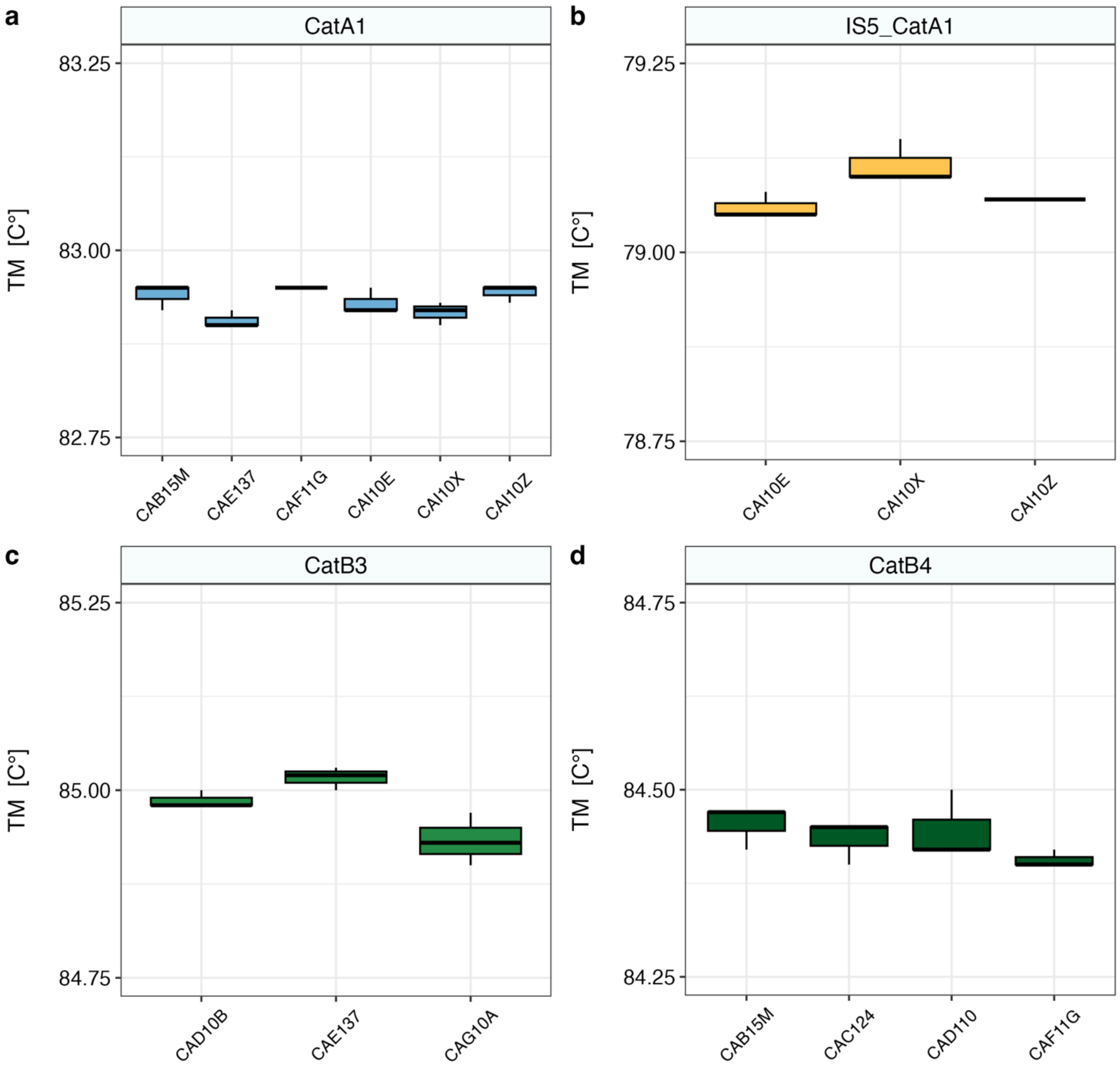


**Supplementary Figure 4 | High-resolution melting (HRM) assay** probing for **a** *catA1*, **b** IS*5_catA1*, **c** *catB3* and **d** *catB4* (= *catB3*$\Delta^{443\text{-}633}$). TM = melting temperature. Each boxplot represents a single isolate with 3 technical replicates.

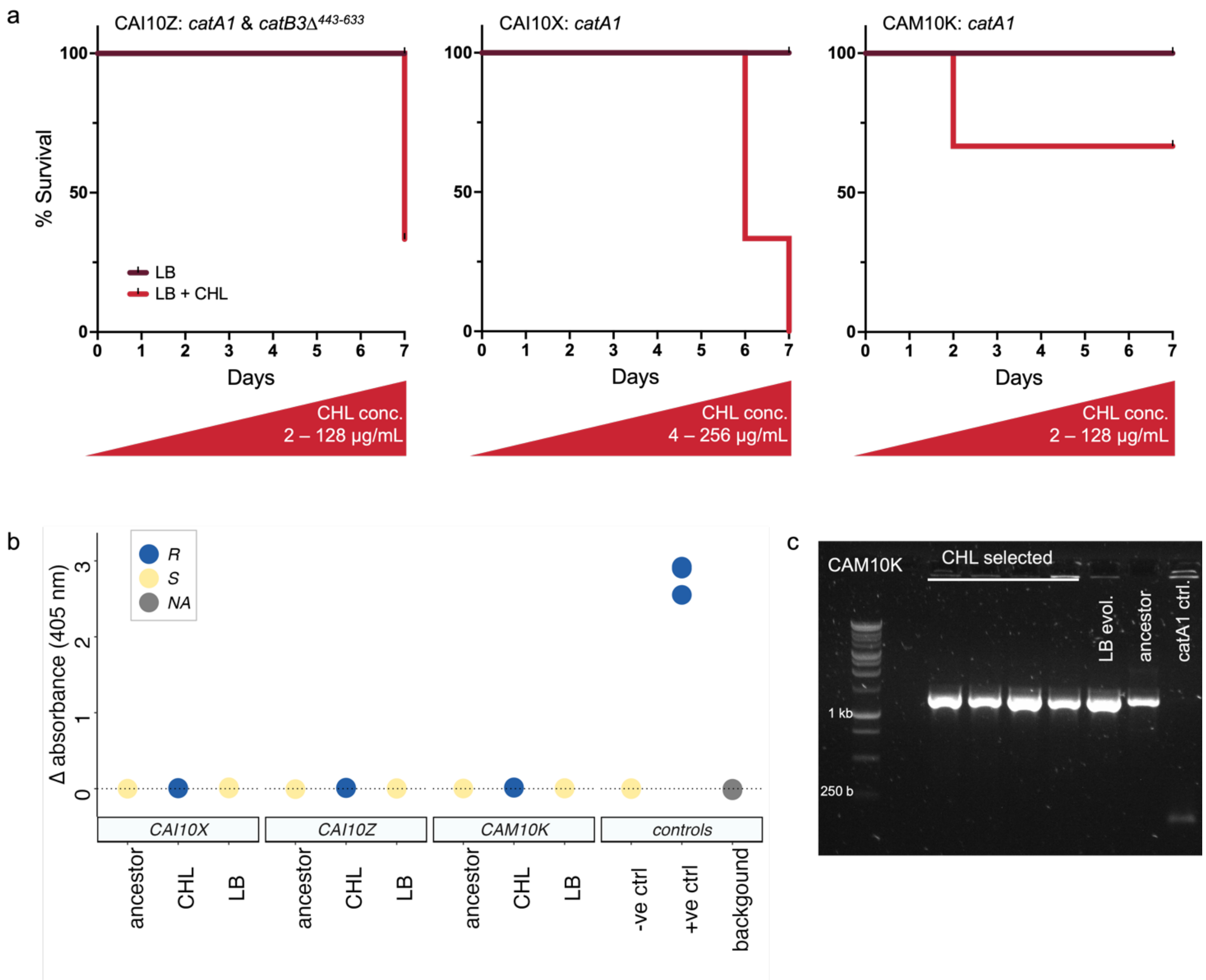


**Supplementary Figure 5 | Stability of IS*5-catA1* upon CHL selection. a** Survival curves of experimentally evolved populations in LB (dark red) and with increasing concentrations of CHL (red) for isolates CAI10X, CAI10Z and CAM10K (all, n=3 for both LB & CHL). **b** rCAT assay performed with last surviving replicate populations of CHL- and LB selected and ancestor population for each isolate. **c** PCR probing for *catA1:* Colony PCR from isolate CAM10K. Lane 1 1kb plus Marker (Promega, UK), lane 2 empty, lanes 3-6 four colonies from 2 independently evolved populations with increasing CHL concentrations from Day 7. Lane 7 LB evolved control population, lane 8 ancestor clone and lane 9 *catA1* wild-type control isolate.

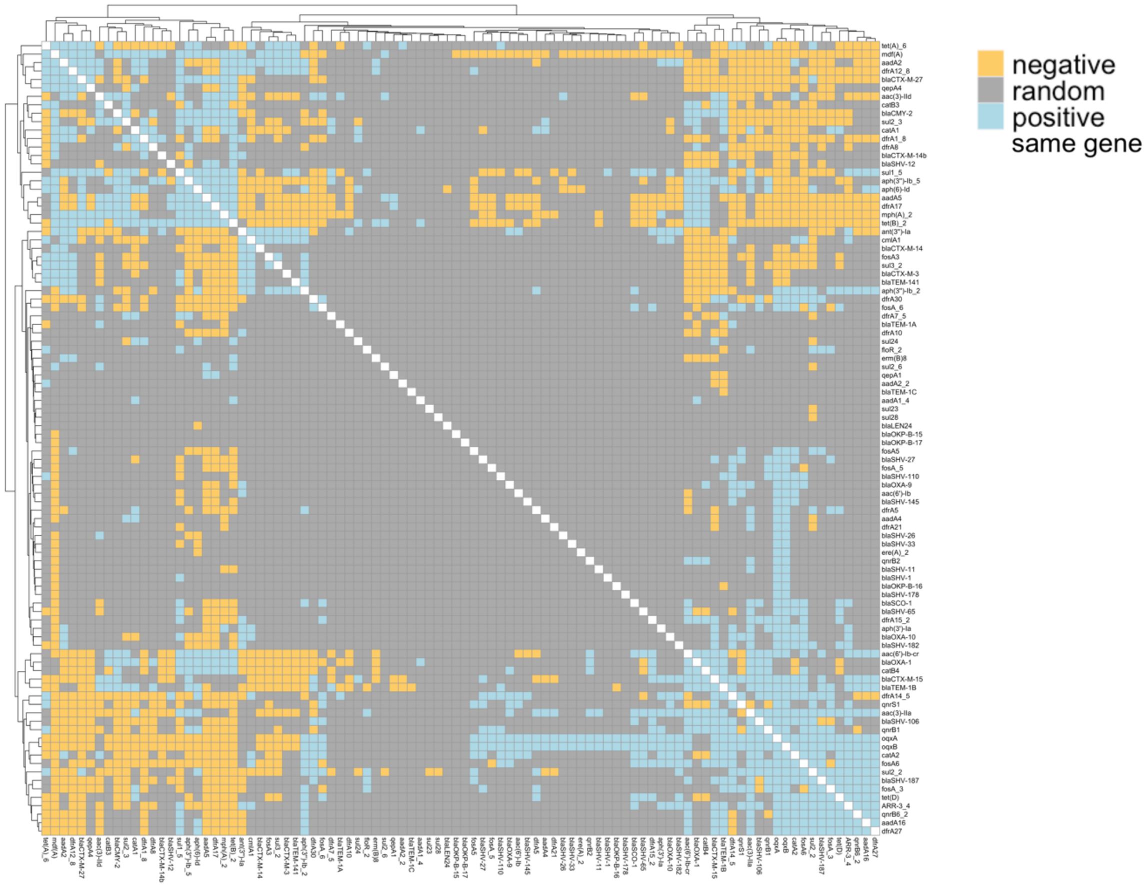


**Supplementary Figure 6 | Co-occurrence networks of AMR genes.** Heatmap displaying co-occurrence relationships between AMR genes as either positive (blue), random (grey) or negative (orange). These are probabilistic values based on the difference in expected and observed frequencies of co-occurrence between each pair of genes, these values were obtained by applying the probabilistic model [1]. Co-occurrence across all AMR genes clustering using Euclidian distance and the complete method

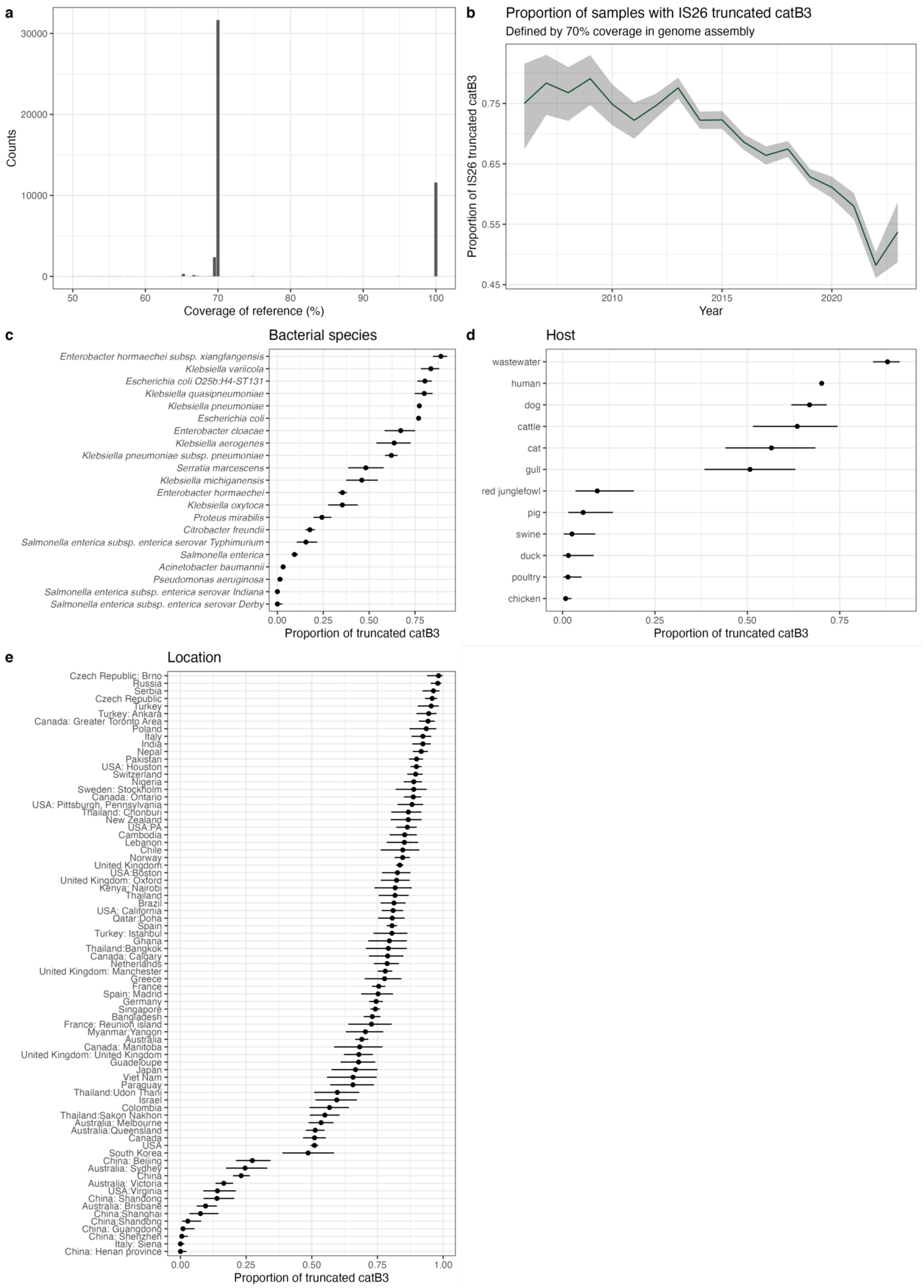
a
Counts
Coverage of reference (%)
b
Proportion of samples with IS26 truncated catB3
Defined by 70% coverage in genome assembly
Proportion of IS26 truncated catB3
Year
c
Bacterial species
Enterobacter hormaechei subsp. xiangfangensis
Klebsiella variicola
Escherichia coli O25b:H4-ST131
Klebsiella quasipneumoniae
Klebsiella pneumoniae
Escherichia coli
Enterobacter cloacae
Klebsiella aerogenes
Klebsiella pneumoniae subsp. pneumoniae
Serratia marcescens
Klebsiella michiganensis
Enterobacter hormaechei
Klebsiella oxytoca
Proteus mirabilis
Citrobacter freundii
Salmonella enterica subsp. enterica serovar Typhimurium
Salmonella enterica
Acinetobacter baumannii
Pseudomonas aeruginosa
Salmonella enterica subsp. enterica serovar Indiana
Salmonella enterica subsp. enterica serovar Derby
Proportion of truncated catB3
d
Host
wastewater
human
dog
cattle
cat
gull
red junglefowl
pig
swine
duck
poultry
chicken
Proportion of truncated catB3
e
Location
Czech Republic: Brno
Russia
Serbia
Czech Republic
Turkey
Turkey: Ankara
Canada: Greater Toronto Area
Poland
Italy
India
Nepal
Pakistan
USA: Houston
Switzerland
Nigeria
Sweden: Stockholm
Canada: Ontario
USA: Pittsburgh, Pennsylvania
Thailand: Chonburi
New Zealand
USA:PA
Cambodia
Lebanon
Chile
Norway
United Kingdom
USA:Boston
United Kingdom: Oxford
Kenya: Nairobi
Thailand
Brazil
USA: California
Qatar:Doha
Spain
Turkey: Istanbul
Ghana
Thailand:Bangkok
Canada: Calgary
Netherlands
United Kingdom: Manchester
Greece
France
Spain: Madrid
Germany
Singapore
Bangladesh
France: Reunion island
Myanmar:Yangon
Australia
Canada: Manitoba
United Kingdom: United Kingdom
Guadeloupe
Japan
Viet Nam
Paraguay
Thailand:Udon Thani
Israel
Colombia
Thailand:Sakon Nakhon
Australia: Melbourne
Australia:Queensland
Canada
USA
South Korea
China: Beijing
Australia: Sydney
China
Australia: Victoria
USA:Virginia
China: Shandong
Australia: Brisbane
China:Shanghai
China:Shandong
China: Guangdong
China: Shenzhen
Italy: Siena
China: Henan province
Proportion of truncated catB3

**Supplementary Figure 7 | Frequency of *catB3* and truncated *catB3*. a** number of isolates (n = 46,667) with *catB3* according to their coverage (in %) to the reference gene. Data downloaded from MicroBIGG-E (4$^{th}$ August 2023). Proportion of truncated *catB3* and confidence interval (**b**) over time including data with > 100 isolates per year, **(c)** by species (> 100 isolates), **(d)** by host (> 50 isolates) and **(e)** geographic location (> 100 isolates). Confidence intervals, where shown, are exact binomial confidence intervals.

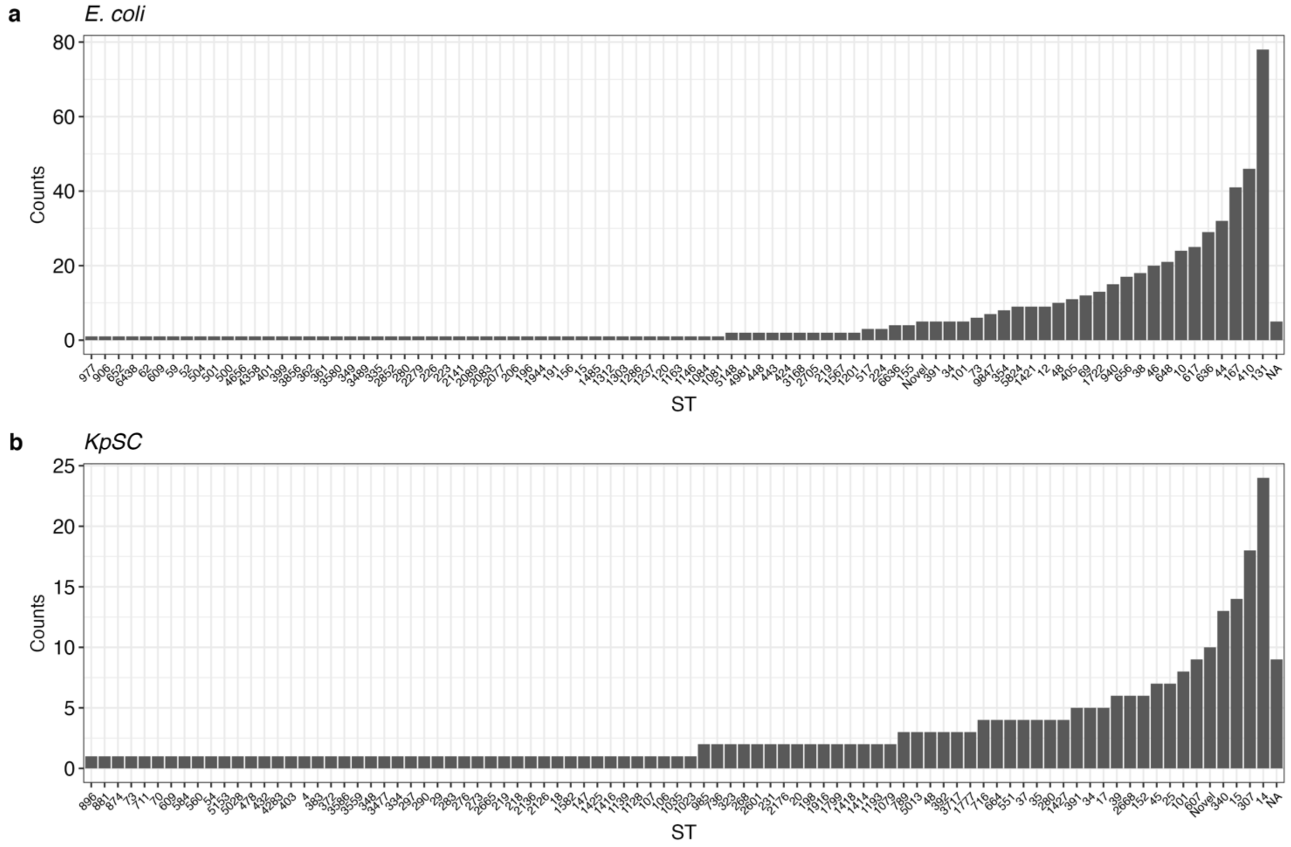


**Supplementary Figure 8: Number of isolates per sequence type** for *E. coli* (a) and *KpSC* (b) of the 840 Malawian genomes included in this study. NA = not determined.

**Supplementary Table 2 – Primers used in this study**

| | Forward Primer (5'-3') | Reverse Primer (5'-3') | Reference |
|---|---|---|---|
| *Cloning primers* | | | |
| catA1-CDS | tttaagaaggagatatacatATGGAGAAAAAAATCACTGGAT | cctgcaggtctggacatttaTTACGCCCCGCCCTGCC | This study |
| catA2-CDS | tttaagaaggagatatacatATGAATTTTACCCGGATTGACC | cctgcaggtctggacatttaTTATTTCAGTATGTTATCACAC | This study |
| catB3-CDS | tttaagaaggagatatacatATGACCAACTACTTTGATAGCC | cctgcaggtctggacatttaTTAGACGGCAAACTCGAGC | This study |
| catB4-CDS | tttaagaaggagatatacatATGACCAACTACTTTGATAGCC | cctgcaggtctggacatttaTTATGGTGATCCCCTGG | This study |
| pEB1-backbone | TAAATGTCCAGACCTGCAGG | ATGTATATCTCCTTCTTAAATCTAG | This study |
| pEB1_sequencing | GCGTATCACGAGGCCCTTTC | AAAGGGAAAACTGTCCATATGCAC | This study |
| *Probing primers* | | | |
| catA1 | AATAAGATCACTACCGGGCGT | GCAACTGACTGAAATGCCTCA | [2] |
| *HRM primers* | | | |
| catA1 | TTTCGTCTCAGCCAATCCCT | CGACATGGAAGCCATCACAA | This study |
| IS5-catA1 | CAGGAACTTGTTCGCACCTT | GGCCGGATAAAACTTGTGCT | This study |
| catB3 | GAAACGCTTCACCGATGAGG | GCCGCTTTGATCTTCTCCAG | This study |
| catB4 | CAATGACGTTTGGATCGGCT | CAACAGTGCCCCACATCTTT | This study |

Lower case = overhangs homologous with pEB1-backbone. NB: Forward cloning primer for *catB3* and *catB4* is identical.

#### Supplementary Table 3 – Long-read assembly statistics, qc and accession numbers.

| ID | Biosample accession number | CHL R/S | CHL-R genes | number of contigs | sum length of all contigs | minimum length contig | average length of contigs | max length contig | Q1 | Q2 | Q3 | N50 |
|---|---|---|---|---|---|---|---|---|---|---|---|---|
| CAE137 | SAMN38266307 | R | catB3, catA1 | 6 | 5,516,907 | 3,204 | 919,484.50 | 5,334,490 | 12,238 | 17,293.50 | 132,388 | 5,334,490 |
| CAD110 | SAMN38266308 | R | catA2, catB3$\Delta^{443\text{-}633}$ | 4 | 5,019,608 | 3,588 | 1,254,902 | 4,867,646 | 10,195 | 74,187 | 2,499,609 | 4,867,646 |
| CAD10B | SAMN38266309 | R | catB3 | 13 | 5,040,963 | 2,972 | 387,766.40 | 4,802,091 | 5,893 | 15,383 | 24,306 | 4,802,091 |
| CAC124 | SAMN38266310 | S | catB3$\Delta^{443\text{-}633}$ | 6 | 5,584,617 | 16,028 | 930,769.50 | 5,315,926 | 16,872 | 29,146.50 | 177,498 | 5,315,926 |
| CAC10A | SAMN38266311 | R | catA1 | 2 | 4,867,656 | 125,633 | 2,433,828 | 4,742,023 | 125,633 | 2,433,828 | 4,742,023 | 4,742,023 |

### Supplementary methods

#### Functional CAT assay (dCAT)

The disk-diffusion CAT (dCAT) assay was adapted from [3]. Cultures of CHL sensitive *E. coli* isolate (EC1010805 from [4]) was prepared in MH2 broth and grown overnight at 37°C, 220rpm. Isolates to be tested and positive and negative control isolates (KI D49363 from [5] and EC1010805) were streaked onto Muller Hinton agar (MH 70191, Merck) and incubated overnight at 37°C. Overnight culture of EC1010805 was diluted 1:10 in MH2 broth and 500 μl spread onto MH agar plates and left to dry for 1 hour at RT. Four discs of 10mm Grade 1 filter paper (1001-6508, Whatman) were added to four points of the agar plate labelled A-D. These related to: A) isolate to be tested with CHL; B) negative control with CHL; C) negative control without CHL; D) positive control with CHL. Using a 1μl loop, a small mass of the relevant bacterial isolate from the MH plate was spread over the filter paper. After a 5-minute incubation at RT, a 30 μg/ml chloramphenicol disc (CT0013B, Oxoid) was placed onto positions A, B and D and a blank disc (CT0998B, Oxoid) was placed onto position C. Plates were incubated overnight at 37°C. QC: Position B should have a zone of inhibition, positions C and D should have no (or a small zone for D) zone of inhibition. If these conditions are met and the isolate of interest (A) does not have a zone of inhibition this indicates CAT activity. If QC checks fail, assay should be repeated.

#### High resolution melting assay

Each HRM assay was performed using 6.25 µL of 2x Type-it HRM PCR buffer (Qiagen, Germany), all primers were added to a final concentration of 400 nM. Molecular grade water was then added to a final reaction volume of 12.5 µl, including 2.5 µl of DNA template. Reactions were thermally cycled in a RGQ 6000 (Qiagen), with the following thermal profile: Taq activation at 95 °C for 5 minutes, followed by 25 cycles of 95 °C for 10 seconds, 58 °C for 30 seconds, 72 °C for 15 seconds. Following this, HRM was carried out by melting from 74 °C to 88 °C, taking a reading in the HRM channel every 0.1 °C, with a 2 second stabilisation between each step. Data was visualised as the negative first derivative of the melting curve to show peak fluorescence dissociation and the predictive Tm of the target was recorded. All analysis was carried out using the RGQ system software. All samples were assayed in triplicate.
